# On the Comparability between Studies in Predictive Ecotoxicology

**DOI:** 10.1101/2025.03.04.641385

**Authors:** Christoph Schür, Kristin Schirmer, Marco Baity-Jesi

## Abstract

Comparability across *in silico* predictive ecotoxicology studies remains a significant challenge, particularly when assessing model performance. In this work, we identify key criteria necessary for meaningful comparison between independent studies: (i) the use of identical datasets that represent the same chemical and/or taxonomic space; (ii) consistent data cleaning procedures; (iii) identical train/test splits; (iv) clearly defined evaluation metrics, as subtle differences—such as alternative formulations of *R*^2^—can lead to misleading discrepancies; and (v) transparent reporting through code and dataset sharing. Our review of recent literature on fish acute toxicity prediction reveals a critical gap: no two studies fully meet these criteria, rendering cross-study comparisons unreliable. This lack of comparability hampers scientific progress in the field. To address this, we advocate for the adoption of benchmark datasets with standardized cleaning protocols, version control, and defined data splits. We further emphasize the importance of precise metric definitions and transparent reporting practices, including code availability and the use of structured reporting or data sheets, to foster reproducibility and advance the discipline.

## INTRODUCTION

The scientific community is making joint efforts to solve grand environmental challenges [1]. This includes those related to chemical pollution and chemical safety. The 350,000 chemicals and mixtures currently globally registered on the market [2] and the practical doubling of chemical industry production since the year 2000, lead to chemical pollution with unknown toxicity to be considered a planetary boundary threat [3–6]. Ensuring the safety of chemicals for humans and the environment is a substantial task that warrants innovative new approaches for hazard assessment outside of classical animal experiments. The legislation for the Registration, Evaluation, Authorisation and Restriction of Chemicals (REACH) by the European Commission requires invertebrate animal tests for chemicals with a manufacturing or import quantity of *≥* 1 tons per year and fish acute toxicity tests *≥* 10 tons per year [7]. Mittal *et al.* estimated the global annual usage of fish and birds to range between 440,000 and 2.2 million individuals at a cost upwards of $39 million *p.a.* [8]. Hence, moving away from the reliance on animal experiments, both ethically and economically, is warranted. Using computational (*in silico*) methods, such as machine learning (ML), to make predictions of ecotoxicological outcomes for chemicals in various species, garners high hopes, but is a yet under-explored approach to environmental hazard assessment [9–11]. Progression of the field of predictive ecotoxicology (and with it the adoption of machine learning methods) will benefit from the scientific community acting collaboratively, with each work building on the findings of the previous [12]. As we will show, this is currently complicated because of a lack of comparability between studies.

Regulatory bodies foremost are committed to ensuring human and environmental safety in line with specific requirements. Ensuring the suitability of alternative methods to augment this system requires a conservative evaluation, leading to cautious progress, not only with the adoption of predictive ecotoxicology methods [13–15]. Much of the literature concerned with predicting ecotoxicological outcomes is focused on quantitative structure-activity relationships (QSAR) models [16, 17]. They are based on the empiric observation that similar compounds will elicit similar biological responses (including toxicity). They have a long-standing tradition in predictive chemistry and can be based on linear and non-linear relationships between chemical descriptors and biological responses. Recent years have seen the adoption of machine learning into QSAR modeling, enabling users to integrate nonchemical data into their predictions [18, 19]. Enhancing a model, beyond the limitation of using only chemical descriptors, ultimately blurs the lines between what can be considered a QSAR model and what goes beyond that. Hence, even though our point ultimately is on the adoption of better practices for comparability in the application of machine learning towards ecotoxicological hazard assessment, it is impossible to exclude QSAR investigations from the scope of this study.

Here, we raise the point that, currently, the field of predictive ecotoxicology is lacking comparability. This, in turn, prevents the transfer of insights between studies and, as a consequence, meaningful progress of the field beyond individual studies. To reinforce this point, we screen a representative subset of available literature on fish acute toxicity prediction and evaluate their comparability based on five comparability criteria that we propose (similarity in dataset, dataset cleaning, splitting, performance metrics, and reporting), outlined in Fig. 1. These criteria are based on similar considerations as *e.g.* the CLEVA-COMPASS proposed by Mundt *et al.* [20] and the EMBRACE checklist by Zhu *et al.*[21]. Although the initial goal of this work consisted of finding the fraction of comparable studies, the collected data led to a concerning conclusion: At the date of data screening, we found no two pairs of studies that fully meet all comparability criteria for ML-based predictive ecotoxicology. Our focus is only on comparability among studies; we do not discuss other fundamental or individual shortcomings that could be the source of other problems (*e.g.* data leakage [22, 23]). This study mainly is aimed at sensitizing the community towards a problem and presenting paths towards improvement.

**FIG. 1.**
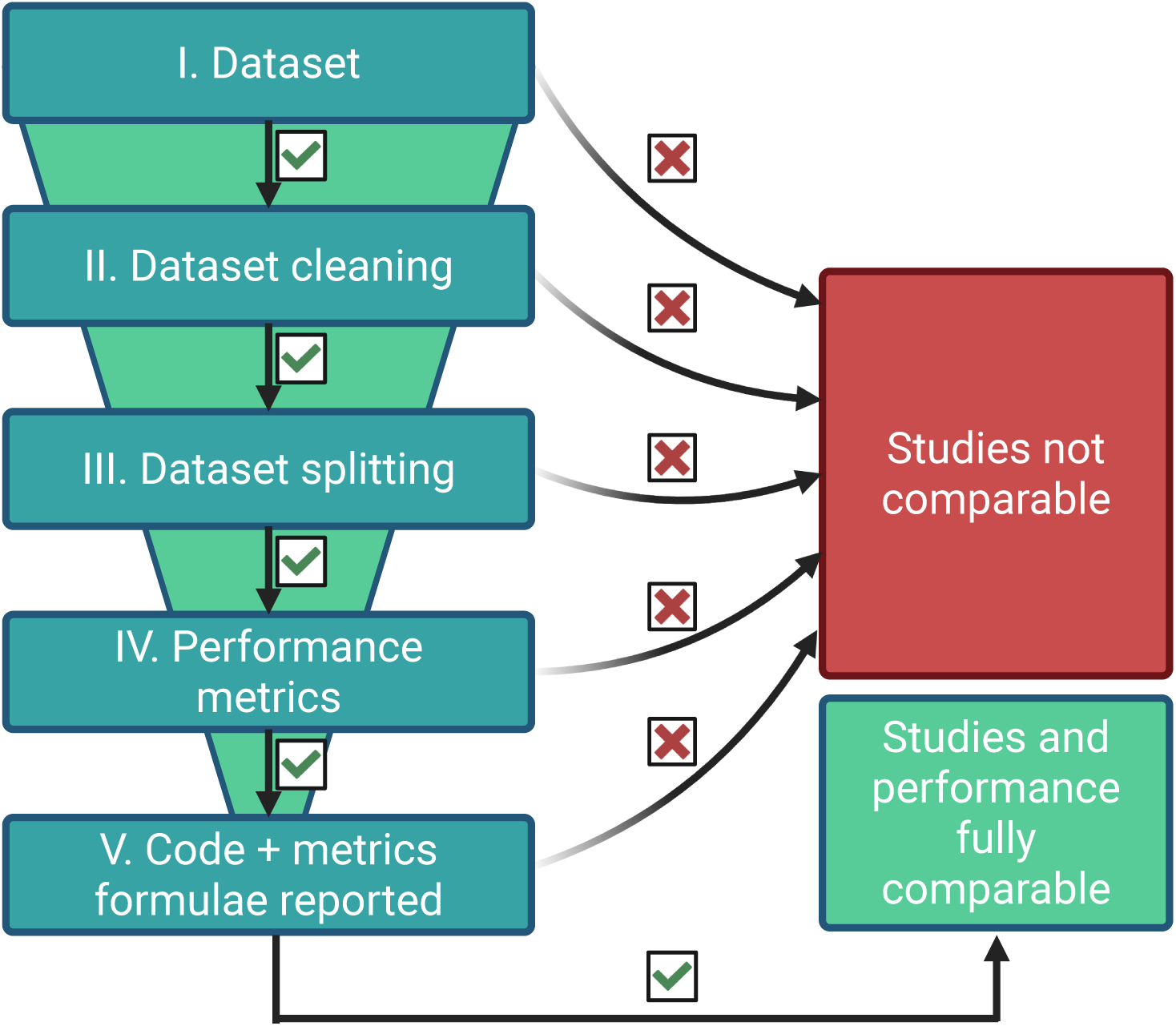
Overview of the comparability levels the studies were judged across. These criteria are elaborated in Secs. I, II, III, IV and V, in the main text. We find that no two currently available studies on *in silico* methods in ecotoxicology meet all of these criteria at the same time.

**FIG. 2.**
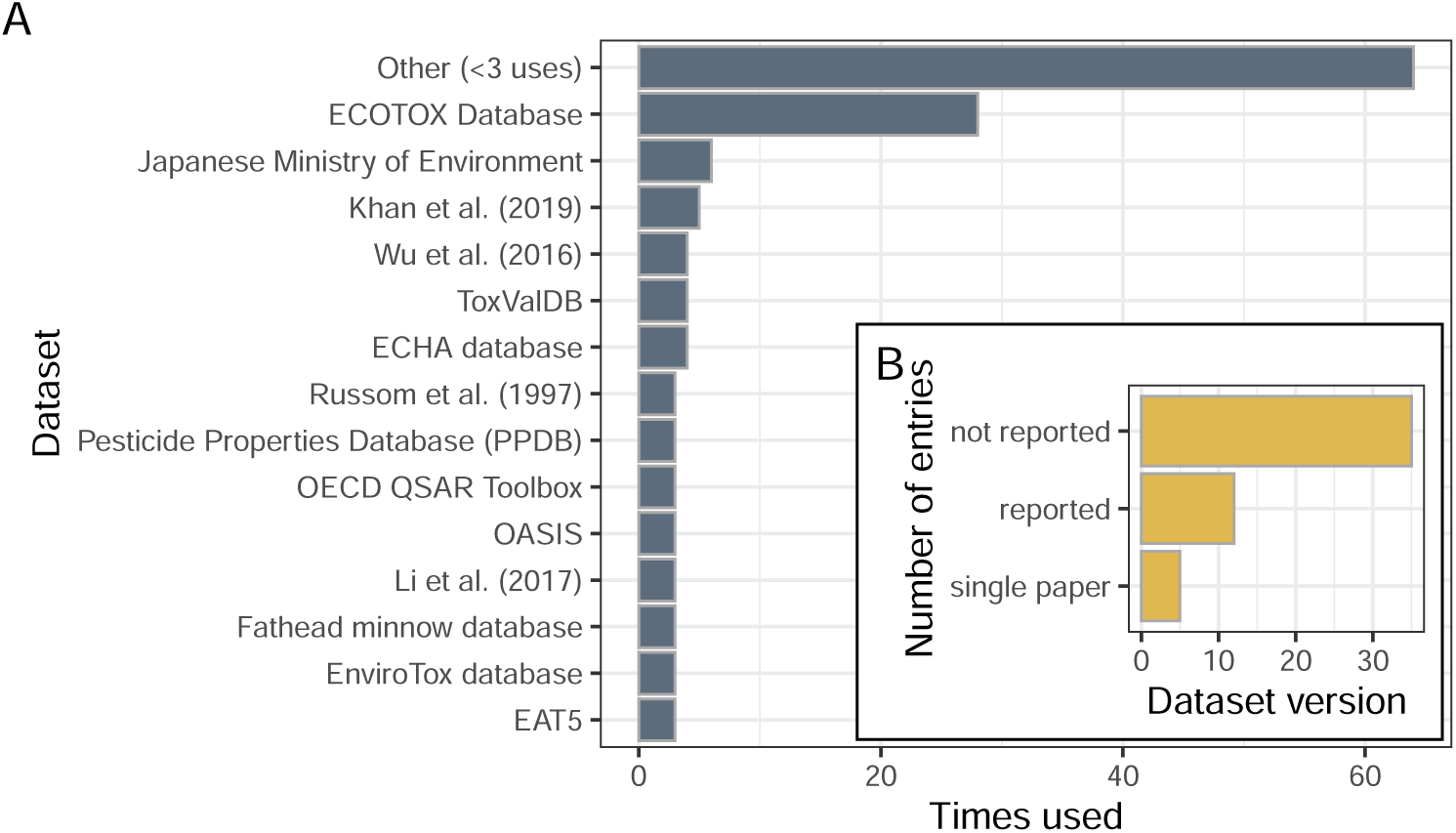
A: Top distinct data sources used across studies in our analysis. Sources used less than three times are aggregated under “Other”; B: Number of times specific dataset versions were reported or single paper sources were used *vs.* not reported.

## MATERIALS & METHODS

### A. Scope & Study selection

We used GoogleScholar with different keyword combinations (*i.e.*, Machine learning, predictive toxicology, ecotoxicology, fish, acute, QSAR), and limited the search results to the period between 2013 and end of 2024. The resulting collection of papers was augmented through adding “similar” and “later” works *via* ResearchRabbit. According the the ResearchRabbit FAQs (accessed 17.06.25), these categories are based on citation networks, assuming that related works cite similar previous works and each other. The resulting dataset of over 100 potentially relevant papers was screened manually and narrowed down to 50 papers that were deemed in scope (*i.e.* removing review papers, studies on other endpoints and taxonomic groups that were brought up through broad use of search terms), representing an overview of the state of the field. Even though the presented work is not a systematic review and not strictly meant to be exhaustive, to our knowledge, it covers the current state of the field and at least the majority of available studies. Studies that only focused on a single group of chemicals were immediately excluded, since it was clear that with such narrow scope there would be no overlap with any of the more broadly-focused studies. Studies could include QSARs making predictions solely based on chemical descriptors as well as studies employing ML methods and/or including a broader range of data types. We settled on acute fish toxicity because it is a vertebrate-based endpoint required by chemical regulation that has been the focus of many predictive ecotoxicology studies, likely due to high data availability and discussions calling for the need of its replacement [24–26]. Additionally, for the purpose of this analysis, the endpoint needed to be clearly defined (*e.g.*, through a test guideline) to try and compare related studies. Obviously, studies concerned with different taxonomic groups (*e.g.* fish *vs.* daphnids) will not be comparable, so increasing the range of studied taxa would not increase the fraction of comparable studies, but rather decrease it. This is why we focus on the narrowly defined scope. Based on these results it is unlikely that broadening the scope would result in a different outcome. The selection criteria do not rule out studies where several taxonomic groups and/or species and/or effect levels (*e.g.*, acute, chronic, reproduction, growth) have been tested in parallel, but our analysis focuses exclusively on the data related to fish acute toxicity.

### B. Data processing and analysis

The information from the selected studies was extracted from the published PDF version of the publications into a table using ChatGPT 4o. The final prompt was engineered through several iterations to yield the most reliable extraction of information across a range of studies and is given in Appendix A. Appendix B contains an example output table. The results were then manually checked for relevance to this work and relevant information. If necessary, the prompt processing was split into parts or additional prompts were given to extract further details or to clarify certain information. Likewise, manual checking of the extracted information against the PDF was performed when results were unclear, and for randomly selected studies as a sanity check. Each study was processed individually in a separate prompt window to avoid confusion between the studies and memory of the model was disabled, *i.e.,* no connection between the individual extractions. The resulting output was manually evaluated, cleaned, and transferred to the table comprising the dataset our study is based on. Further cleaning, summarizing, and visualization of the data was conducted in R (Version 4.4.1) with RStudio (Version 2024.09.0+375). The code and data of the analysis is available on https://github.com/mbaityje/COMPARABILITY.

### Limitations

We acknowledge that the extraction of (sometimes complex) information using a large language model may produce errors. This is particularly likely in the case of the definitions of performance metrics, because they appear to not always be recognizable by the model, especially when not explicitly referred to in the text or embedded as images. The same is true where information is not phrased in an easily comprehensive way or implicitly. Manual checking of a subset of extracted information revealed that the observed error rate was comparable to manual data extraction, but at vastly higher throughput. Nonetheless, even when expecting a certain error rate and thus consider our analysis more qualitative than strictly quantitative, the emerging patterns remain. This notion is based on the fact that comparability already is not given on the level of datasets and cleaning.

As for the number of considered studies, this could have been increased by extending our analysis to studies considering other taxonomic groups (*e.g.*, other common ecotoxicological test organisms, such as crustaceans or algae) or effects and endpoints. However, within ML applied to ecological hazard assessment, fish mortality is the most used test, *i.e.* the one with the highest probability of finding comparable studies. Thus, extending our work to less used effects or taxonomic groups would likely not change the conclusions.

## RESULTS & DISCUSSION

### Comparability criteria

We propose the five levels of comparability based on two branches of reasoning: I. factors that influence model performance, II. factors that enable or prevent reproducibility of published modeling results. The former category approaches the problem from the side of technical considerations towards applying ML models towards the same problem and where studies could differ, leading to different and incomparable outcomes. The second category is mindful of the scenario of a researcher wanting to fully replicate a modeling approach. The criteria are thus born from practical experience as well as theoretical considerations. For full comparability between two studies, all criteria should be met. At the same time this ensures maximal transparency and reproducibility.

#### I. Datasets

To support comparability, and thus enable comparison of model performances independently of different training data, researchers should use the same datasets. This criterion comes with the caveat that what ultimately matters for comparability is the applicability domain (AD) of the dataset, *i.e.* the kind of species and chemicals, the range of chemical properties, and outcome variables (toxicity) that is represented [27]. One cannot expect models to achieve quantitatively comparable performances across datasets that, for example, contain different chemical groups or only partially (if at all) overlapping ranges of molecular weight, octanol-water partition coefficient (*LogK_ow_* or *LogP*) or other descriptors. The same is true for data based on differing experimental designs, *i.e.* not derived from the same standardized test guideline, such as the OECD test guideline 203 for fish acute toxicity [28]. Fixed benchmark datasets are already commonly used in other branches of ML-based research to exclude variability in used datasets as a driver of model performances and to ensure comparability [29–32]. The 50 studies we include in our analysis contained relevant toxicity data originating from 82 sources (Fig. B-A shows the datasets that were used at least three times). Many of these data sources, such as the US EPA ECOTOX database (accessed: 18.06.2025), which was the most commonly used single source (Figure B-A), are updated frequently and are thus not a static resource that can be linked to without indicating the access date/version of the dataset. The dataset source is more clearly defined when it is associated with a single scientific paper, which was only the case in 10% of cases. More often, datasets from different sources were combined, resulting in a new dataset (even when all sources were defined individually). In this case, the final dataset a study is based on heavily depends on data cleaning. For the majority of the screened studies (*>* 70%), the concrete version of the used dataset was not reported (Figure B-B).

### Different datasets for different purposes

Different datasets are needed in order to answer different research questions. Benchmark datasets primarily serve the purpose of allowing to compare model performances on similar grounds. Additionally, they can provide a baseline model performance if the goal is to gauge the impact of additional features on model performance. For this, a model is trained and tested on the benchmark dataset before it is expanded with the features to be studied and trained and tested again. Likewise, models trained on different datasets can be benchmarked against a benchmark dataset to give a comparable reference of performance and to test against a (potentially out-of-domain) external dataset. This is the more realistically feasible option opposed to replicating several published models based on the provided code and test them on one’s own dataset. In order to answer different scientific domain questions, different benchmarks need to be created and published within the community. As part of *ADORE*, we published several specific challenges focusing on a number of research interests, such as training exclusively on crustacean and algae data to predict mortality in fish or training on single species data to predict on several species. More broadly, Ashauer and Jager (2018) proposed a concerted effort of the community to fill data gaps for combinations of chemical classes, modes of action, and taxonomic groups to achieve full data coverage across these levels and reduce bias [33]. For taxonomic coverage, something similar is currently underway within the Precision-Tox initiative [34]. The process to create datasets for specific purposes and with high-stakes decisions, such as regulatory relevance, in mind can be guided by documents, such as the “Datasheets for datasets” [35].

##### Take home message

Studies aiming for broader contextual relevance may benefit from aligning with commonly used datasets to be comparable with previous studies. If no datasets for this purpose exist yet, they should be created.

#### II. Cleaning & aggregation

Data cleaning (selecting or removing specific data points, dealing with duplicates, outliers, missing values, standardization of the data) affects the dataset composition and applicability domain and, thus, the questions a researcher is able to ask and the scope of the answer a model is able to give. Potential reasoning underlying these decisions can range from purely domain-specific, like limiting data to a range of a chemical property or species found in a certain geographic region, to considerations stemming from technical limitations (compatibility of data with model types, constraints imposed by molecular representations). Aggregation of multiple values into a single value per species, chemicals, or experiments (depending on the data structure and purpose of the study), can be an integral step of data cleaning.

Often, cleaning is not reported transparently enough for the steps to be fully reproducible (Figure 3). This would include a list of all filtering and removal steps that were employed and would also be aided by code availability (see Sec. B). Frequently, it is not even clear if data was aggregated using the geometric or arithmetic mean (Figure 3-A). Studies can only be considered comparable if the dataset and cleaning match and are reported thoroughly, ideally with accompanying code (see section 6). We acknowledge that not all datasets require extensive cleaning, especially those originating from single papers (around 10 % of the studies screened here) or that do not contain duplicates or missing values. Authors also may deliberately choose to not remove outliers or normalize the data, which is then not explicitly mentioned in the paper. These cases are, hence, also included into the “not reported” group.

**FIG. 3.**
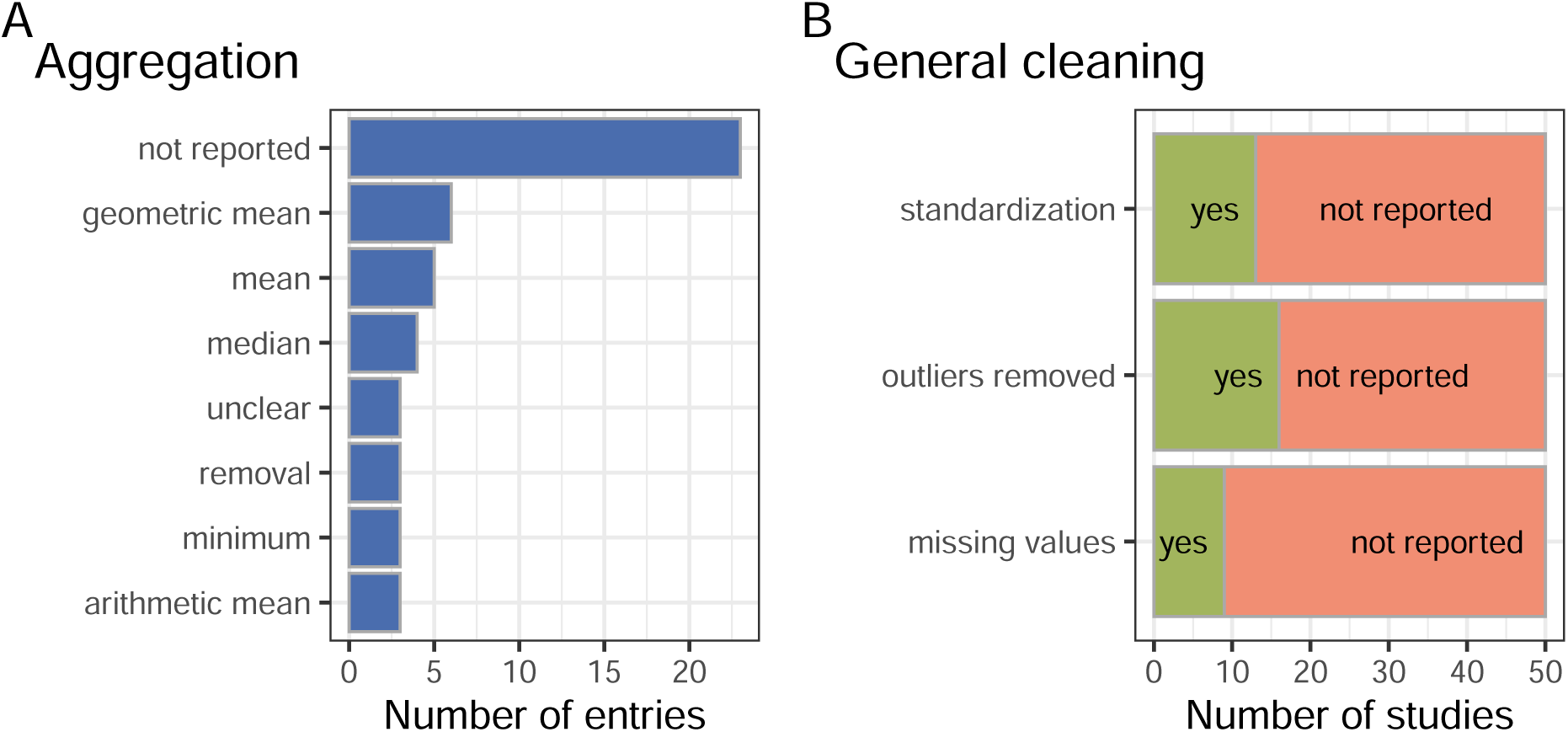
Reporting of important cleaning steps across studies: A: Aggregation of duplicate values; B: Standardization of data points, treatment of missing values, and removal of outlier values.

###### Take home message

Data set version and cleaning must be the same in order to allow for comparability.

Transparent documentation of the cleaning process and applied criteria and filters, ideally supported by providing scripts, enhances reproducibility.

#### III. Splitting

Splitting is the process of dividing a dataset into the training, validation, and test set and is an integral part of assessing model performance. Model training and hyperparameter-tuning is performed on the training+validation set, while the test set is kept completely separate until the final phase of the study where model performance is put to the test on it. Several ways to achieve a train-test-split are available: Random splitting, stratified random splitting, or splitting according to other aspects of the data, like chemical scaffolds [36] (*i.e.* a shared molecular backbone of molecules). Random splitting is the most commonly used approach (Figure 4-A), and is done by randomly distributing data points between the train and test sets at a pre-determined ratio (often 80 % of the data points go to train and 20 % to test; Figure 4-B). Stratification by toxicity (”Stratification, toxicity”) is the second most common strategy, while stratification by chemicals was used less often. “Not applicable” for the splitting ratio was applied when an external test set was used that did not require splitting of the dataset. As it was already highlighted in previous work, the way the data are split can artificially inflate performances [37, 38], so it is crucial to ensure that splittings are kept under control. More practically, we show this in our previous work on *ADORE*, where random splitting (with a high likelihood of data leakage) led to a root mean square error (RMSE) of 0.5, which corresponds to the highest achievable performance based on the original variability of the training data. Meanwhile, splitting in a way that the train and test set contain different chemicals, ergo asking the model to make a prediction on chemicals it has never seen, decreases the performance to an RMSE of around 0.9-1 [39]. The effect of splitting was more profound than the use of molecular descriptors. The danger of data leakage has recently been pointed out by Stock *et al.* (2023) for ecological applications of machine learning [23]. One approach to circumvent this issue, as described in our example from Gasser *et al.* (2024), is a stratified split according to the chemical (and ideally also by species, if several species are included in the dataset), so that all data points related to each chemical are put into either the train or the test set (analogous reasoning is true for the validation set). This is still not ideal, since similar chemicals can elicit a similar toxicological response (notwithstanding activity cliffs, chemicals that have a very similar structure but highly different biological responses [40]). Here, splitting by the molecular scaffold, *i.e.* the basic backbone structure, can be an option [36]. This substantiates the importance of data splitting for assessing and comparing model performances.

**FIG. 4.**
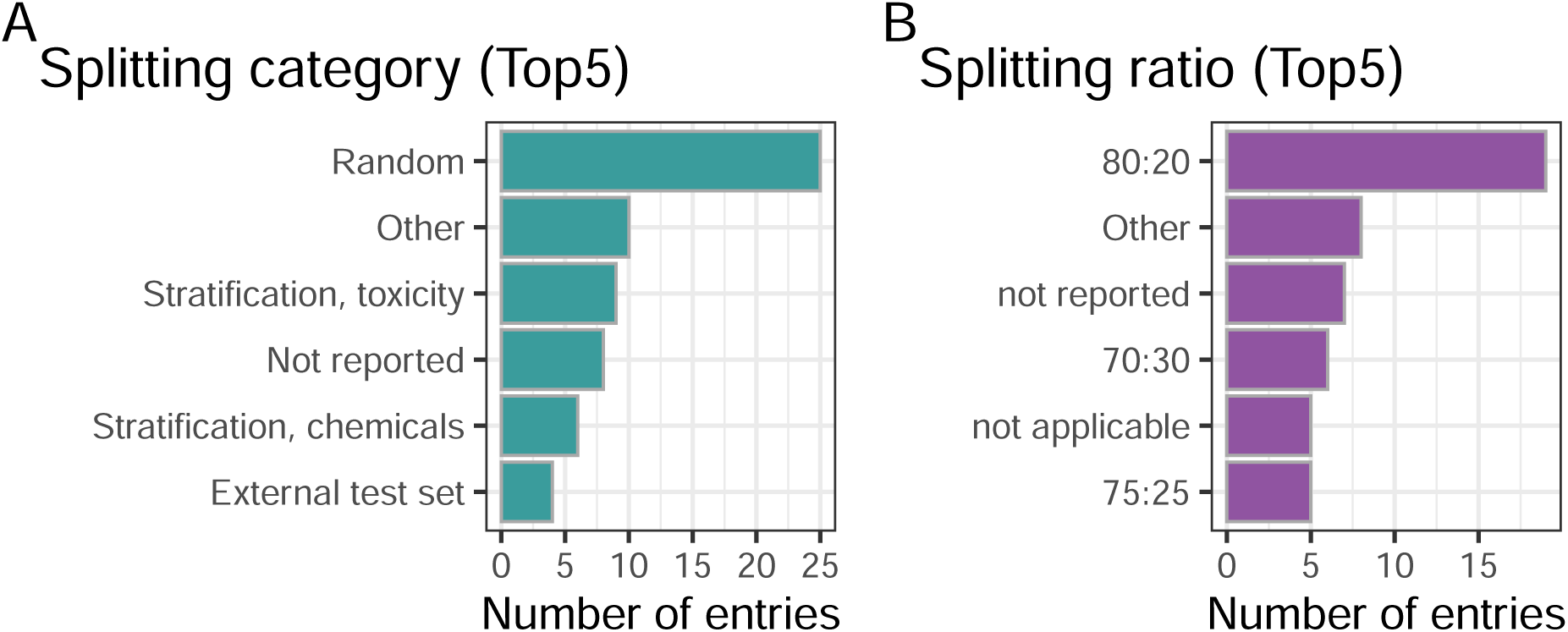
Overview of the way in which training and testing dataset splitting was achieved (A) and the most often reported splitting ratios (B).

Finally, we highlight that it is not sufficient that the splitting strategies are the same: the exact data entries in each of the splits should be the same. As an idealized example, let us take two works, A and B, performing a 1:1 train/test random splitting in a dataset where 50% of the examples are easy, and 50% are hard. If work A has all the easy examples in the training set, and work B has them in the test set, the two works will report different performances (B will arguably show higher performance), even if both A and B are using the same exact method. Therefore, if we want the performances in the two works to carry the same meaning, the exact data instances appearing in each of the data splittings should be the same. This implies that if a study wants to be comparable to future work, it has to report the exact composition of training and test set (while there is some flexibility with the validation set).

###### Take home message

Between studies, even with the same exact dataset, reliable comparisons require consistent and clearly documented data splitting strategies.

#### IV. Metrics

Model performance is assessed through performance metrics, depending on the modeling task and usually on a hold-out test set after training a model on the training set. Several kinds of metrics are commonly used, even though their naming and definition is not always made clear within the studies. Figure 5-A highlights the most commonly reported performance metrics (metrics that were not part of the 10 most common are summarized under “Other”). Reported designations for similar metrics differed between studies, hence they were manually summarized under broad umbrella terms. Sensible performance metrics differ between modeling tasks, *i.e.* regression modeling *vs.* classification. The most common task was regression followed by binary classification, followed by classification with different numbers of classes (Figure 5-B). Lastly, we checked how many of the studies defined their metrics, either directly through equations or by referencing a specific function from a software package that was used to calculate (Figure 5-C). Here, roughly 50% of the studies did not adequately define the performance metrics. Concurrently, only stating the name of a metric is often not sufficient to adequately identify them, as we argue in the following.

**FIG. 5.**
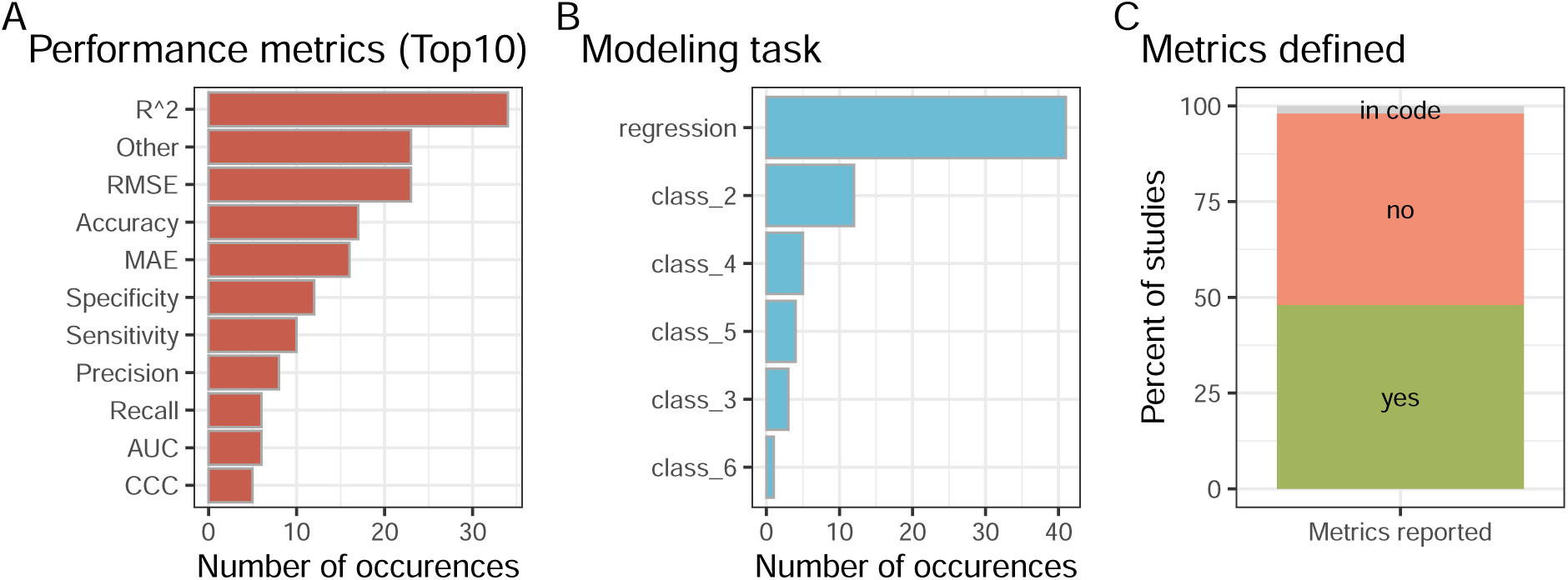
A: Ten most commonly reported performance metrics (less-commonly used metrics are summarized under “Other”); B: Reported modeling task (class n refers to classification task with n number of classes); C: Are the metrics clearly defined/reported, either through equations or by referencing specific functions from software packages. RMSE = Root Mean Square Error; MAE = Mean Absolute Error; AUC = Area Under the Curve; CCC = Concordance Correlation Coefficient.

##### Performance metrics

We now highlight that metrics that appear to be the same can sometimes be different. Therefore, one needs to 1) accurately report the formula used to calculate the performances and 2) possibly adhere to the same exact measures. As an example, we will take the *R*^2^ metric (often called *coefficient of determination*), which is both the most widely used (see Fig. 5–A), and also problematic as far as comparability is concerned [41–44]. The general expression of the *R*^2^ is

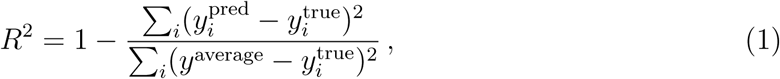

where *y*_i_^pred^ are the model predictions about each test data point *i*, *y*_i_^true^ are the ground truths (or measured values), and *y*^average^ is the average value of the ground truths. If the *R*^2^ is close to 1, the predictions are good, if it is around 0, the predictions are equally as good as taking random points around the average. If they are negative, the model is worse than just taking the average.

The *R*^2^ metric simplifies in different ways for specific kinds of models, usually depending on whether these have intercepts or whether they are generalized linear models [41, 44]. For this reason, some software and communities that are focused on specific kinds of models, may use specific definitions of the *R*^2^ which are equivalent to Eq. (1) only under specific conditions. One example is that, for linear problems, the *R*^2^ reduces to the squared Pearson correlation coefficient, *r*^2^ [45, 46]. Ref. [41] collects previous literature and shows that 8 different definitions of *R*^2^ are used. Through some simple examples, it further shows that each of these metrics can give rise to different values. This multiplicity of ways to calculate the *R*^2^ can lead to conflicts in the results, when they are used under conditions that do not guarantee their equivalence to Eq. (1).

A further ambiguity arises when splitting in train/test sets [44].^1^ Which data split is *y*^average^ evaluated on? Training set, test set, or entire dataset? Ref. [44] states that Eq. (1) “should all relate to test data, not training data”, but it also recognizes that others advocate for calculating *y*^average^ on the training set. From the point of view of comparability, all strategies are acceptable, as long as it is clearly stated how *y*^average^ was calculated, and comparisons are only performed among studies which adopt the same definition. We also highlight that the Mean Square Error 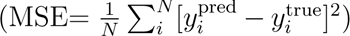 measures similar deviations as the *R*^2^, without requiring to calculate *y*^average^, thus being less ambiguous. However, as we will see in the next section, even the MSE, as any other performance metric, can be ambiguously defined.

###### Take home message

The exact definition of the used metric should be provided. Some metrics depend on the splittings too, so their definition should also be specified in relation with the splittings.

##### Relevance of repeated entries, micro and macro averaging for performance metrics

Ecotoxicological experiments are often performed more than once. While, in many cases, one can aggregate these values by *e.g.* taking the median or geometric mean, it is often desirable not to perform any aggregation. This is for example the case of Bayesian approaches, where it is assumed that the output does not necessarily converge to a single value, but it could rather only be well-represented through a distribution of points [18]. Regardless of the reason, if repeated entries are not aggregated, one must be especially careful to, besides not including the same entry in both training and test set, specify how the given metric deals with repeated entries.

As an example, let us take a metric where every single datapoint has the same weight (these are usually called micro-averaged, in opposition to macro-averaged metrics where sets of datapoints are first aggregated according to some rule to not over represent a certain chemical, species, or experiment). If 50% of the points in the test set are related to glyphosate tested on one or more species, then models with good performance on glyphosate will be advantaged with respect to others. This can be undesirable, since it may be by pure chance that glyphosate appears in the test set. It can also be desirable, since perhaps we want to use our model in a geographical region where glyphosate is very present. In general, when micro-averaged metrics are used, since those are agnostic about repetitions, every repetition in the test set gives more weight to that chemical. This is equivalent to making the hidden assumption that the chemicals that are tested more often are more relevant for applications, which is not always reasonable. One example of different ways to define a metric which is apparently always the same appears in Ref. [18], where four different definitions of the root mean square error (RMSE) are explicitly provided. These different definitions of the RMSE (micro-averaged, micro average chemicals and macro average taxa, macro average chemicals and micro average taxa, macro-averaging taxa and chemicals) represent different desires at the level of interpretation of the model performance.

###### Take home message

The metrics should take repeated entries into account, by explicitly specifying how they are dealt with.

#### V. Providing the code and dataset

Since some metrics can be calculated in different ways and metric names are not always consistent or unique, it is possible that the software used to implement metrics may not align with the intended metric definition for a given use case. The best way to allow for comparability in this case is to allow people to recalculate the metrics themselves. This is achieved through increased transparency, *i.e.*, making FAIR (Findable, Accessible, Interoperable, Reusable) models with all the code associated with a publication available [12] (this could include all the code used for model training and tuning (if applicable) in addition to the fully trained model). Code availability was recently highlighted by Gundersen *et al.* (2024) to be the most important factor towards successful replication of a ML-based study, irrespective of the quality of documentation of said code [47]. Likewise, Cronin *et al.* (2023) argue that following the FAIR principle could positively affect regulatory acceptance of models [12]. Figure 6-A shows the fraction of screened papers where the corresponding code was made available. Fewer than 50% of studies make their code freely accessible. Reproducibility of published work requires the use of the exact dataset, hence the value provided by freely available benchmark datasets that are published independently of associated works [48]. Figure 6-B indicates that roughly 60 % of the screened papers made their data available or linked to an unambiguous source where the dataset was available. However, if given, the data are often not provided in an easily machine-readable format (*e.g.*, comma-separated values (*csv*)) but rather as a table as part of the supplementary information (often in a word document) that requires further extraction/cleaning. In addition to the code, the field would greatly benefit from adopting universal reporting standards, such as the “Model cards for data reporting” [49], REFORMS [50], and EMBRACE [21].

**FIG. 6.**
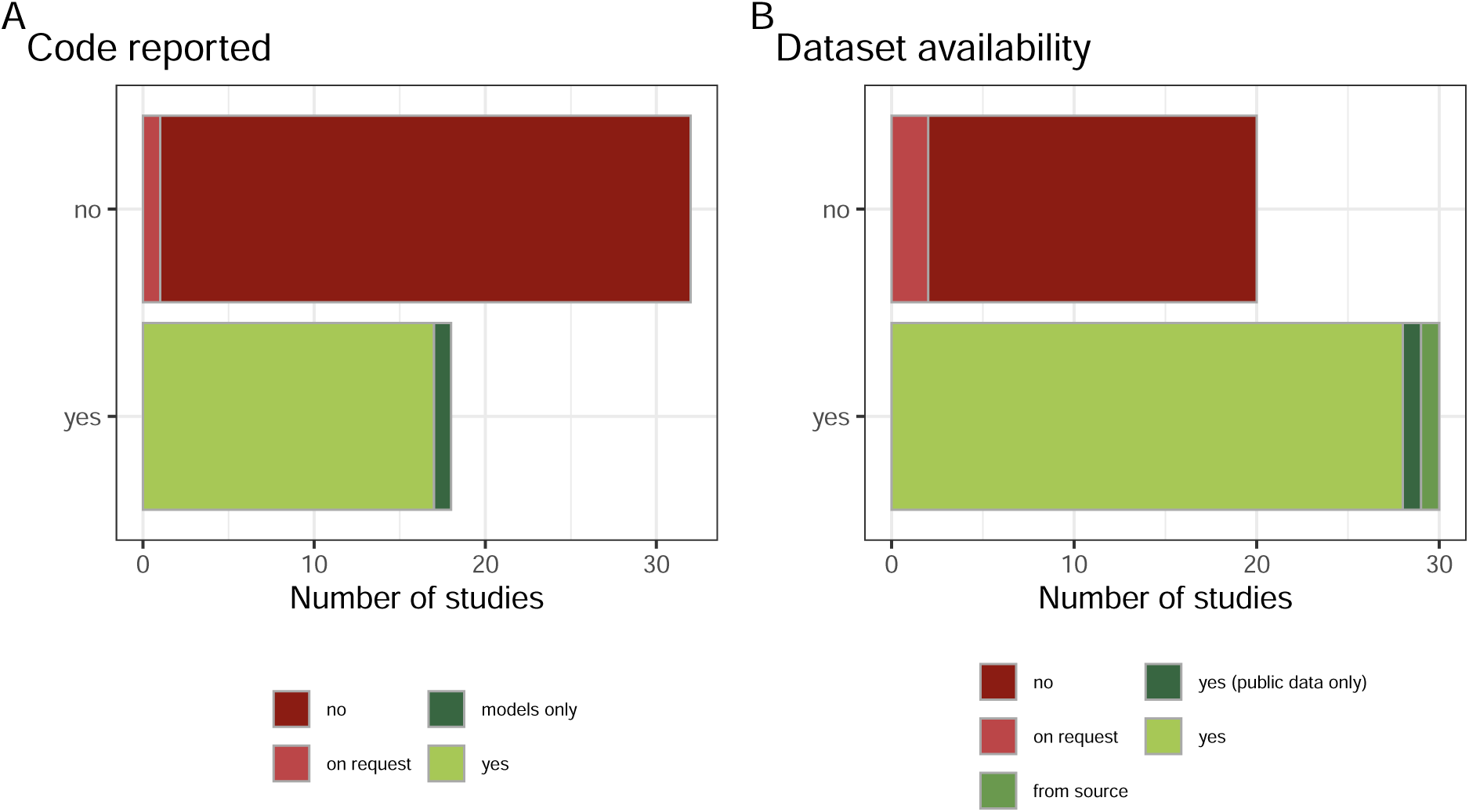
A: Number of studies that made their code available; B: Number of studies that made their dataset available.

###### Take home message

Making data and code publicly available can significantly improve reproducibility and comparability.

Data sheets with all the necessary information should be easily accessible.

## EXAMPLE COMPARISON

Up until this point our discussion of comparability between studies have been abstract and conceptual. Here, we want to highlight a tangible example to show how easily comparability can get lost. The two studies by Ghosh *et al.* [51] and Yu and Zeng [52] both use the toxicity data published by Li *et al.* [53]. We select these two studies as an example, because they use a dataset from a single paper, which removes most ambiguity associated with the use of databases or combined sources (data selection, different versions *etc.*) that may not easily be deciphered. Such a dataset can be considered fixed, at least on the level of toxicity data. Researchers employing such dataset still are left to perform their own cleaning (if desired/necessary), additional feature curation, and to perform the train-testsplit. Hence, both studies start out with the same dataset containing 1258 pesticides. Both remove inorganic compounds and metals, while Ghosh *et al.* further manually removed “unknown chemical structures, salts containing organic polyatomic counter ions, mixtures, and substances of unknown or variable composition”. These data cleaning steps are, for the most part, caused by data gaps for chemical descriptors and technical restrictions of the software used to calculate them. Already at this point the studies cannot be compared to each other, even though they started out using the exact same data. Furthermore, they move on to train models for regression (Ghosh *et al.*) and binary classification (Yu and Zeng), split their data 75:25 (method unclear) and 80:20 (based on euclidean distance of chemical descriptors), respectively. Moreover, Yu and Zeng use the whole dataset containing multiple species, while Ghosh *et al.* train different models for rainbow trout, bluegill sunfish, and “misc species”. We summarize the comparison between these two studies in Tab. I. We emphasize once again that we are not criticizing these researchers and their work, but rather showing a tangible example of two studies being based on the same toxicity data without the produced models being comparable due to the processing steps. Other studies already diverge at the very initial step of the data source.

**TABLE I.**
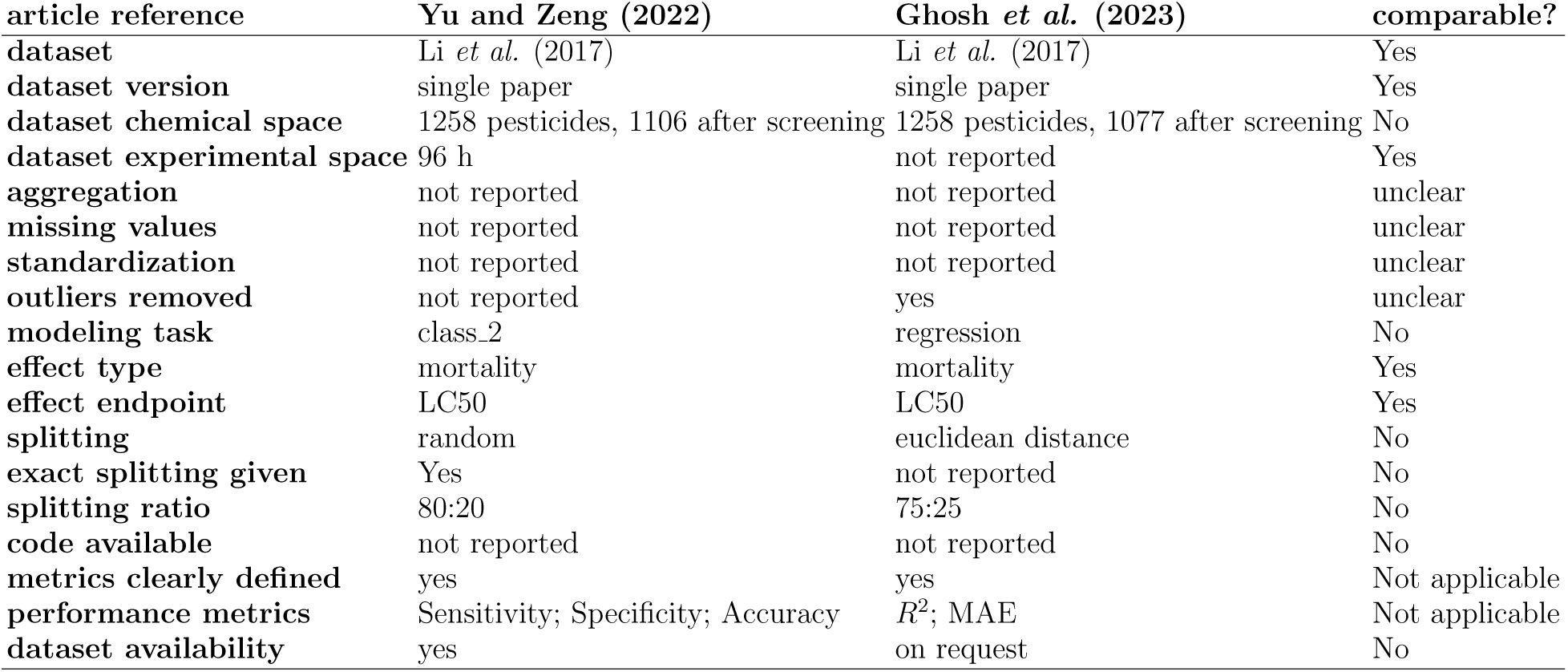
Exemplary comparison of two studies: Yu and Zeng (2022) [52] and Ghosh *et al.* (2023) [51], initially based on the exact same dataset (Li *et al.* (2017) [53]) and how their comparability can be judged based on given information.

### SUGGESTIONS TOWARDS ACHIEVING COMPARABILITY

Machine-learning-based research in general faces fundamental issues in terms of implementation, reproducibility, and reporting [12, 22, 23, 54–58]. Several researchers addressed these shortcomings and suggested checklist-based solutions to harmonize model implementation and reporting [12, 21, 59–61] and the adoption of benchmarks for the sake of comparability [29–32]. These issues are not unique to any field of ML-based research, but different branches have been quicker or slower in the adoption of best practices. In the field of predictive ecotoxicology, we see that progress is already hindered by the lack of comparability among available studies. Model performances can only reasonably be compared to each other when they were obtained on the same dataset, covering a similar range of species and chemical space. This can be addressed through the use of *benchmark datasets*: curated, freely accessible, and commonly accepted datasets that can form the basis of comparability. Benchmark datasets constituted the basis for AI revolutions in diverse fields, from computer vision [62–64] to hydrology [65, 66], and are arguably a fundamental element needed for an AI revolution in predictive ecotoxicology [29, 67].

Such datasets include documented and explained cleaning, fixed splitting *etc.*, and more importantly, they are “frozen”, meaning that everyone using them trains and tests their models in the exact same conditions. Performances on these datasets can then be tracked through centralized documentation databases, such as Papers with Code: *“a free and open resource with Machine Learning papers, code, datasets, methods and evaluation tables”* (https://paperswithcode.com/about, accessed: 16.06.2025). Once typical performances on a specific benchmark data set are established, it becomes straightforward to know how good performances need to be in order to be considered competitive against other. However, out of the over 12,000 datasets listed on Papers with Code as of June 2025 (accessed: 18.06.2025), 69 are labeled as “biology”, highlighting the large potential for growth.

Standardized datasets at the same time solve several fundamental problems for different stakeholders: Authors spend less time curating and cleaning data as well as documenting these steps. In turn, authors from diverse background can more readily apply their ML expertise to challenges in (potentially understudied) fields (think: ecotoxicology *vs.* human-centered toxicology). Documentation is further streamlined through the use of standardized reporting sheets, such as REFORMS [61] and EMBRACE [21], and adhering to best practices for the implementation of machine learning and QSARs into ecotoxicology [12, 59, 68, 69]. This also reduces potential frustration about the time investment to find out specific information on model development or the data from a study, if it is given at all. This enables the interested audience to immediately know where to find the desired information for a study and other researchers to know how their model performs compared to others. Oftentimes, such relevant information is difficult to locate without structured reporting. As editors and reviewers, members of the community can ask researchers to adhere to comparability and reporting standards, in turn reducing time devoted to review work. Special attention needs to be given to definitions and reporting of performance metrics, ideally in concert with code documentation. Communally working on improving the standards of model development and transparency will ultimately progress the field as a whole, getting ML closer to helping progress environmental hazard assessment in ways many are already envisioning it to do.

To provide a solution to this need within the realm of predictive ecotoxicology and to reduce the barrier of entry for researchers lacking the biological/ecotoxicological expertise to compile, clean, and contextualize such data themselves, it is necessary to use datasets that serve as a benchmark for machine learning in ecotoxicology. To our knowledge, two datasets with such stated purpose exist: the ADORE dataset [70] and the dataset by Svedberg *et al.* [71]. In addition, the EnviroTox database [72] is a curated ecotoxicology dataset based, among others, on the ECOTOX database (accessed: 18.06.2025) that could serve this purpose. However, for many applications it would need further cleaning to be suitable for specific questions. These datasets should serve as a point for comparison between methods addressing similar questions. However, it is good practice to use more than one benchmark dataset. First, different benchmark datasets might be suited to address different kinds of research questions. Different modeling architectures require different feature sets (*e.g.*, regression modeling based on many features [18] *vs.* recommender systems based on few [73]). It would then be insightful to assess whether a method fulfills specific modeling needs, leading towards the refinement of the best suitable methods. Second, the usage of benchmark datasets implies that the test set is always the same throughout several studies. However, there is a risk that new methods become tailored to perform best on a specific benchmark dataset [74], and thus may superficially be judged as useful, while it fails in terms of generalizability beyond this specific benchmark set. In the case of large language models (LLMs), where benchmarking is not as straightforward, attempts have been made to achieve better comparison of model performances through platforms such as LMArena (accessed 17.06.26). To overcome this, new benchmark datasets should be proposed every few years in order to avoid that the developed methods are too influenced by the specific choice of a dataset.

Regardless of the scope of the individual datasets, ML learning studies should report their results on as many benchmark datasets as possible to demonstrate generalizability of the model performance. This proves the robustness of the results, and allows to track the progress of the state of the art. Furthermore, the usage of benchmark datasets must be complemented with a precise reporting of the used performance metrics, and reproducible code.

## CONCLUSION

Our analysis of current and past literature on the prediction of acute ecotoxicological outcomes in fish highlights that the usual way of describing and reporting modeling limits comparability of model performances on equal grounds. This, however, is a prerequisite to finding out which methods are the most effective, irrespective of different underlying datasets. To address this, we propose a list of criteria through which one can assess if two studies are comparable to each other to encourage the community to be mindful of factors influencing comparability. Our examination spans key factors such as dataset selection, modeling tasks, and performance metrics, emphasizing the need for greater consistency and transparency. Our findings indicate that no existing studies fully meet these criteria, underscoring a persistent gap in the field. While these concerns may not be new and have been raised in several other branches of ML-based research, they continue to hinder broader adoption and integration of methods — something we highlight through a representative bibliographic analysis of studies aimed at the same ecotoxicological outcome of fish acute mortality. We believe that establishing clearer standards for comparability, aided for example by the use of defined and fixed benchmark datasets and reporting standards, is a necessary step for advancing machine learning applications in predictive (eco)toxicology and fostering collaborative progress in the field.

## ACKNOWLEDGMENTS

We are thankful to L. Gasser from the Swiss Data Science Center (SDSC) for inspiring conversations. This work was made possible through the SDSC grant “Enhancing Toxicological Testing through Machine Learning” (project NoC20-04) and partly carried out in the framework of the European Partnership for the Assessment of Risks from Chemicals (PARC; Grant Agreement No. 101057014). Figure 1 was created with BioRender.com.

## AUTHOR CONTRIBUTIONS

CS: Conceptualization, Data curation, Formal analysis, Investigation, Methodology, Software, Validation, Visualization, Writing – original draft; KS: Conceptualization, Funding acquisition, Project administration, Supervision, Writing – review and editing; MBJ: Conceptualization, Data curation, Formal analysis, Funding acquisition, Investigation, Methodology, Project administration, Software, Supervision, Validation, Writing – original draft. All authors reviewed and approved the final manuscript.

## DECLARATION OF GENERATIVE AI AND AI-ASSISTED TECHNOLOGIES IN THE WRITING PROCESS

During the preparation of this work the authors used *ChatGPT 4o* in order to extract information from published PDF versions of the papers screened for analysis. Further information is given in section B. *ChatGPT 4o* was used for language improvements. After using this tool, the authors reviewed and edited the content as needed and take full responsibility for the content of the publication. No generative AI or AI-assisted technology was used for generating the text of this manuscript.

## Footnotes

1 Our argument also applies to validation splitting, but for the sake of clarity we omit it without loss of generality.

## Notes

https://github.com/mbaityje/COMPARABILITY

### Competing Interest Statement

The authors have declared no competing interest.

### Summary of Updates

The revision was carried out based on the feedback of two anonymous reviewers at the journal Computational Toxicology. Several new paragraphs have been added and over 100 changes have been made to the manuscript.

