## Supplementary Material for "On the Comparability between Studies in Predictive Ecotoxicology"

### Appendix A: ChatGPT prompt

The following prompt was given to ChatGPT to extract the desired information from the PDF version of the studies.

```
Please answer a couple of questions about this scientific article.
    Please do so based on the whole document, not only parts. Do
    it step by step and make the output a table. Include the
    authors, paper title, publication year, and the doi in the
    table. Please answer each of the questions step by step. If
    nothing related to the question is included in the article
    state ``not reported".

What is the source of the toxicity data? Is there information
    about which database version was used or the date of retrieval
    for the database? Was there more than one data source that was
    used (a combination of several databases or scientific papers
    as sources)? Please specifically list each of the sources
    individually.

What kinds of chemicals are included in the dataset? This could be
     something like organic chemicals, pesticides or some other
    description of the chemical range.

What kind of species or taxonomic groups are included in the
    dataset? How many different species are in the dataset? Was
    there some specific selection criterium that was followed?

Was there a selection of specific experimental parameters, like
    limiting the data to include only specific experimental
    durations (e.g., 24 hours, 48 hours etc.)?
```

Was the data aggregated in any way? Example here include calculation of mean or medians for multiple values or just keeping multiple values as data points.

Was there any mention of missing values and how they were handled? This could include removing them or some form of imputation. If this is the case, please quote the text mentioning this.

Was the data standardized in any way before use? This could mean that it was reduced to minimum or maximum values. If this is the case, please quote the text mentioning this.

Is there a mention of outliers and how they were handled (for example removed according to a specific criterion)?

What was the modeling task? Classification, regression or both? If it was classification, how many classes were used?

What was the effect type that was modeled? This could be something like mortality, lethality, immobilization, intoxication, growth, behavioral alterations etc.

What was the endpoint that was modeled? This could be something like lethal concentration 50 (LC50), effective concentration (EC50), no observed effect concentration (NOEC), lowest observed effect concentration (LOEC).

When splitting the data points in training, validation and test set, did the authors separate the data in groups based on the values of some target variable or were they distributed randomly? Splitting can be by random, by chemical, by species or by some other parameter.

What was the ratio of splitting between either train/training and test/testing data? Was there a validation set as well? 80:20 is very common, but other ratios are possible.

Is the code for the performed modeling available, either on github, on request, through a repository, in the supplementary information or not at all?

What re the performance metrices used to evaluate model performance? This could be something like R^2, r^2 (note that there is a difference between a metric with an upper-case and a lower-case r), RMSE, MAE, Accuracy, F1 Score, Specificity, Sensitivity. There can be multiple different ones at the same time.

Are equations/formulas given for how to calculate the performance metrics or are the metrics otherwise defined?

Is the dataset used in the study provided through a repository or as part of the supplementary information?

Please summarize all the answers to the questions in a table.

### Appendix B: Example ChatGPT output

This section presents an example of the output ChatGPT produced from the prompt in Sec. A for the PDF of Gasser et al. (2024).

| Question | Answer |
|---|---|
| **Authors** | Lilian Gasser, Christoph Schür, Fernando Perez-Cruz, Kristin Schirmer, Marco Baity-Jesi |
| **Paper Title** | Machine learning-based prediction of fish acute mortality: implementation, interpretation, and regulatory relevance |
| **Publication Year** | 2024 |
| **DOI** | 10.1039/d4va00072b |
| **What is the source of the toxicity data?** | ADORE dataset based on the ECOTOX database, harmonized and pre-filtered. Additional data from various sources were included. No version/date was reported. |
| **What kinds of chemicals are included in the dataset?** | Primarily organic chemicals. |
| **What kind of species or taxonomic groups are included?** | Fish (140 species), crustaceans, and algae. No specific species selection criteria were elaborated. |
| **Was there a selection of specific experimental parameters?** | Yes, data includes specific durations: 24, 48, 72, and 96 hours, transformed for modeling. |
| **Was the data aggregated in any way?** | The dataset was filtered and harmonized, but individual data points were retained without averaging across experiments. |
| **Was there any mention of missing values and how they were handled?** | Not explicitly mentioned. |
| **Was the data standardized before use? If yes, please quote the text.** | Yes, continuous features were standardized based on the training data. Quote: "The continuous variables were standardized based on the training data." |

| | |
|---|---|
| **Is there a mention of outliers and how they were handled?** | Not explicitly mentioned. |
| **What was the modeling task? Classification, regression, or both?** | Regression. |
| **What was the effect type that was modeled?** | Mortality. |
| **What was the endpoint that was modeled?** | Lethal concentration 50 (LC50). |
| **When splitting data into training, validation, and test sets, was the separation random or based on criteria?** | Two splits: random sampling (totally random) and stratification by chemical compound (split by occurrence of chemical compounds). |
| **What was the ratio of splitting between training and test data? Was there a validation set?** | Training and test sets followed an 83:17 ratio. Validation was conducted using 5-fold cross-validation on the training data. |
| **Is the code for the performed modeling available?** | Yes, the code is available through a public repository. Quote: "All code was written in Python 3 using established ML libraries. It is available in a public repository." |
| **What performance metrics were used to evaluate the model performance?** | RMSE (root mean square error), MAE (mean absolute error), $R^2$ (coefficient of determination). |
| **Are equations/formulas given for how to calculate the performance metrics?** | Yes, equations for metrics like RMSE, MAE, and $R^2$ are provided in the appendix. |

| | |
|---|---|
| **Is the dataset used in the study provided through a repository or supplementary information?** | The dataset (ADORE) is a benchmark dataset, but no specific link to a repository was mentioned for direct access in this paper. |
